# Acute Smurf mortality and age-dependence in a two-phase ageing model: Statistical inference from drosophila and mice experimental data

**DOI:** 10.64898/2026.02.18.706552

**Authors:** Luce Breuil, Marie Doumic, Sarah Kaakaï, Michaël Rera

## Abstract

Ageing is traditionally conceived as a continuous process of progressive physiological decline. However, recent evidence across species suggests that ageing may instead proceed through distinct phases. Using state-of-the-art statistical methods, we develop a rigorous analysis of longitudinal survival data from 1,159 individually tracked female *Drosophila melanogaster*. This data-driven analysis leads us to introduce a new parametric model of transition rates within the two-phase ageing framework. Flies were monitored using the Smurf assay, which detects increased intestinal permeability through leakage of an ingested blue dye, and is a strong biological marker of ageing. The Smurf phenotype identifies a sharp transition from a non-Smurf state to a Smurf state that precedes death. Our results yield three key findings. First, the Smurf transition rate follows a Gompertz-Makeham law, increasing exponentially with age. Second, contrary to previous constant-rate assumptions, newly transitioned Smurf flies exhibit remarkably high mortality - approximately 40% die within 24 hours - followed by an exponential decline in death rate that stabilises to a lower constant baseline. Third, we identified a mild but statistically significant negative dependence between time spent non-Smurf and subsequent Smurf lifespan. Our best-fit model captures a potential bimodal nature of mortality curves using simple, biologically interpretable functions. Validation using data from two mouse strains confirms the broader applicability of this framework. These results establish a quantitative foundation for the two-phase ageing paradigm and highlight a critical period of vulnerability immediately following the physiological transition to frailty.

## Introduction

Ageing still lacks a consensual definition^1^. There is a common underlying notion of time passing, that nevertheless affects living organisms in a broad range of modalities^2^. It is mostly considered as a continuous process characterized by a progressive alteration of molecular and physiological functions, ultimately leading to a time-dependent increasing mortality risk^3^.

This conception of the ageing process strongly impacts, and is impacted by, experimental designs. In the laboratory, we mostly study ageing using synchronized cohorts of individuals compared across distinct chronological time-points. An implicit assumption is that each of these cohorts are homogeneous, individuals within constitute biological replicates. The idea that ageing is not a linear process affecting individuals in a progressive way brings novel views on that aspect.

The first mention of a biphasic ageing process that we found in the literature dates back from 1961 where JM. Clarke and J. Maynard Smith concluded “that the life span of a fly can be divided into two phases, referred to as ‘ageing’ and ‘dying’“^4^. Both phases have distinct physiological properties, resistance to environmental stressors and mortality trajectories.

In 2011, Rera and colleagues presented for the first time the Smurf phenotype, an age-dependent abrupt increase of intestinal permeability to a small blue food dye - hence the name of the phenotype. Its age-dependent incidence can be modulated by lifespan-increasing intervention^5^. This simple *in vivo* assay allowed for the identification of two subpopulations of individuals in any given *D. melanogaster* population. A first, non-Smurf subpopulation, and the Smurf one composed of individuals showing an extended blue coloration due to increased intestinal permeability. They later showed that the rate of non-Smurf to Smurf transition increases as a function of time at a rate negatively correlated with life expectancy. That is, it reflects biological age more than chronological age. Smurf individuals show physiological changes usually described in the literature as hallmarks of ageing^6^, including loss of energy stores, altered motility, or increased inflammation. These changes actually encompass all^7^ the transcriptional hallmarks of ageing^8^, which are a quantifiable form of the broadly used hallmarks of ageing^3,9^. We also showed the Smurf transition to be a necessary phase of life^6,10^, although other groups have suggested otherwise under different experimental protocoles^11–13^.

More importantly, this biphasic process for ageing is broadly evolutionarily conserved as it was described in nematodes, drosophila or zebrafish^14^, and more recently in mice^15^. Across all these organisms, the Smurf transition precedes death from natural causes with a scaling between organisms^16^.

Recently, Shen and colleagues^17^ showed, using multi-omics data, that ageing in humans progresses in distinct waves rather than continuously. In demographics, this ‘unexplained’ heterogeneity across ageing humans is often modeled by latent variables called frailty. Multiple parameters allow the identification of frail individuals from high-level behaviours such as grip strength and walking speed^18^ down to low-level molecular indicators such as circulating citrate or very-low-density lipoprotein particle size^19^. They allow a 5-year mortality prediction with hazard ratios ranging from 0.65 up to 3.5. In addition, intestinal permeability to the same blue food dye we are using in flies has been shown to predict high-risk of mortality in critically ill^20^ and septic patients^21^.

The broad evolutionary conservation of this two-phase ageing process, as well as its stereotyped molecular characteristics, led us to introduce a two-phase model of ageing for synchronised populations^10^.

Based on an initial analysis of experimental observations^6,10^, the proportion of Smurfs over time was first approximated by a linear equation^6^. This simple assumption allows the modelling of Drosophila longevity curves using experimental measurement. This simple assumption allows the modelling of Drosophila longevity curves using experimental measurement. However, this should be viewed as a first-order approximation, as it does not incorporate the full biological complexity or take into account all the mathematical constraints of the models.

The transition rate from the Smurf state to death was rather well described by a constant, by fitting the experimental survival curve of Smurf individuals with an exponential function e^-kt^, with k the death rate. This equation was obtained by fitting experimental survival times of Smurfs from 5 different genetic backgrounds and multiple ages. Furthermore, the descriptive analysis of the experimental data did not indicate a clear relationship between the duration of the Smurf phase and that of the preceding non-Smurf phase.

In the past decade, the Smurf phenotype was validated in other model organisms for ageing including nematodes, zebrafish and mice. The estimation of Smurf survival in these different organisms suggests a scaling of the Smurf duration with the life expectancy at birth of the considered organism. That is, while Drosophila Smurfs survive for a median time of 2-3 days, mice survive 2-3 weeks and zebrafish approximately 6 months. An important question is then to assess how the model developed using fly data can actually be transferred to other organisms.

The aim of this paper is to introduce a rigorous mathematical framework and to take advantage of recent developments in statistics, in order to challenge and complement results previously obtained in Tricoire and Rera^10^. Our approach is guided by a systematic investigation of the data, based on as few a priori assumptions as possible. In this spirit, we introduce a general two-phases compartmental model of ageing, in which individuals transition from a non-Smurf compartment to a Smurf compartment before death. A rigorous non-parametric statistical analysis provides the basis for developing a biologically relevant parametric model of the transition rates and the potential dependence between the durations of the two phases.

Using a rigorous non-parametric statistical framework^22^ applied to individual lifetimes of *D. melanogaster*, we establish four principal findings. First, the rate of non-Smurf to Smurf transition follows a Gompertz-Makeham law, revealing that this discrete physiological switch accelerates exponentially with age rather than occurring at random. Second, and most strikingly, approximately 40% of flies die within the first 24 hours of turning Smurf, after which the death rate declines sharply. This critical window of acute vulnerability was previously overlooked under constant death rate assumptions. Third, we uncover a mild negative dependence between the time spent in each phase: flies that remain non-Smurf longer die faster once they transition, consistent with continuous sub-threshold damage accumulation operating within the non-Smurf phase. Finally, cross-species validation on two mouse strains suggests that this two-phase ageing process is conserved beyond Drosophila. The Gompertz-Makeham Smurf transition rate and the high early post-transition mortality are both recovered in mice, establishing the basis for a broadly applicable tool for ageing research.

## Results

### 1. Model and non-parametric analysis

These last few years have seen a strong interest in the development of a discontinuous model of ageing. This conceptual change highlights the need for a better understanding of the mathematical structure underlying survival, and raises several questions, such as: How many phases are relevant to describe ageing? Which equations best describe these phases? Previous work^10^ was able to model longevity curves using two phases under some approximations. The present study aims at assessing thoroughly which mathematical models of ageing capture the main features of the data using a rigorous data-driven statistical approach.

We rely on data published in Tricoire and Rera^10^, consisting of the individual observation of m=1159 mated female flies, from age 11 days to their death, with a life expectancy at age 11 of around 20 days (480 hours). In the following, for time spent non-Smurf and total lifetimes, time 0 corresponds to day 11, age at transfer on blue food. In brief, flies maintained in individual vials throughout their lives were scored daily for survival, death and the Smurf phenotype. This experiment showed that flies can die only in the Smurf stage: no non-accidental death in the non-Smurf stage was reported. On average, the time spent in the non-Smurf phase (phase 1) is around 16 days post transfer on blue food (387 hours) while the time spent in the Smurf phase (phase 2, physiologically old individuals) is about 4 days (93 hours).

Let us describe the setting. The total lifetime of each fly is denoted by *τ*_*i*_, As schematised in Figure 1, *τ*_*i*_ can be separated into two consecutive phases, non-Smurf and Smurf, of respective duration 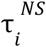 and 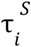 such that 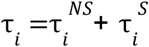. The distribution of each of these variables can be characterised by their hazard rates. The hazard rate k of a random event occurring at time *τ* is the instantaneous rate at which the event can occur (see Supplementary Information).

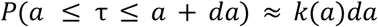

**Figure 1.**
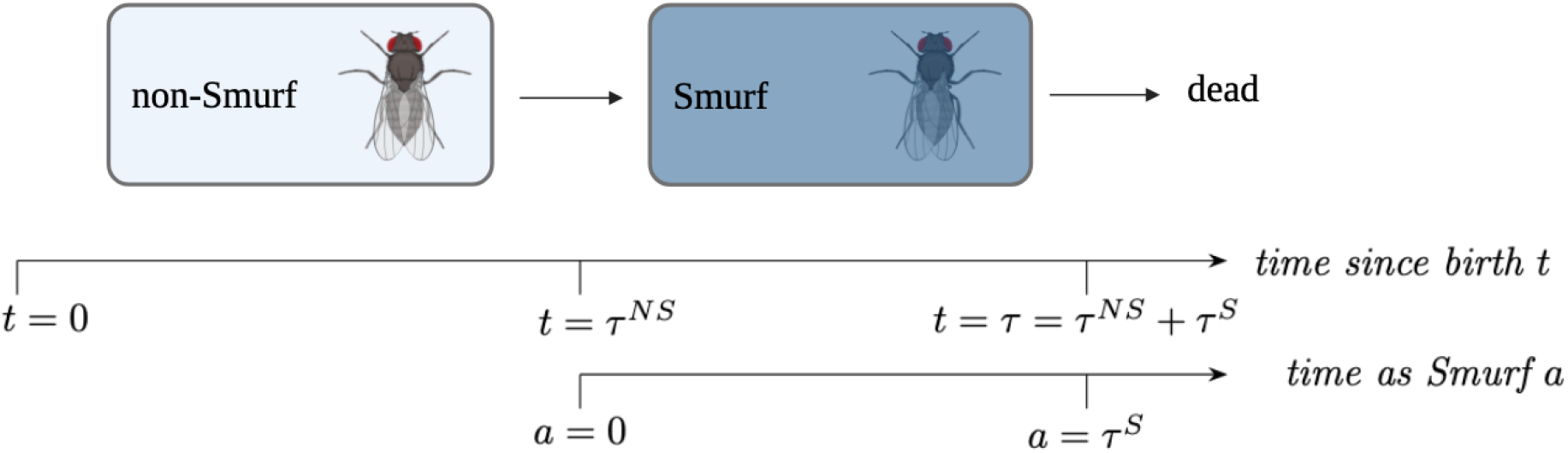
Schematic representation of the two phases of ageing.

The hazard rate of *τ*_*i*_ corresponds to the population-level mortality rate which is classically estimated, and is called apparent death rate in the following. On the other hand, the hazard rate of 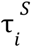 is the actual death rate, as it characterises the transition from life to death. Finally, the hazard of 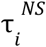, which is the transition rate from non-Smurf to Smurf, is called Smurf transition rate in the following.

In Figure 2, we present the three survival curves (grey dots) and hazard rates (red curves) associated respectively with the distribution of the total lifetime (Figure 2A), the time spent in the non-Smurf state (Figure 2B) and the time spent in the Smurf state (Figure 2C). To avoid imposing prior assumptions, we rely on a data-driven non-parametric approach based on recent statistical results on the estimation of hazard rate functions^22^ (see also Supplementary methods). The 95% bootstrap variability estimation error band (in grey in Figure 2), together with the Q–Q plot Figures S1 and S2, indicate a good fit of the hazard rate estimation.

**Figure 2.**
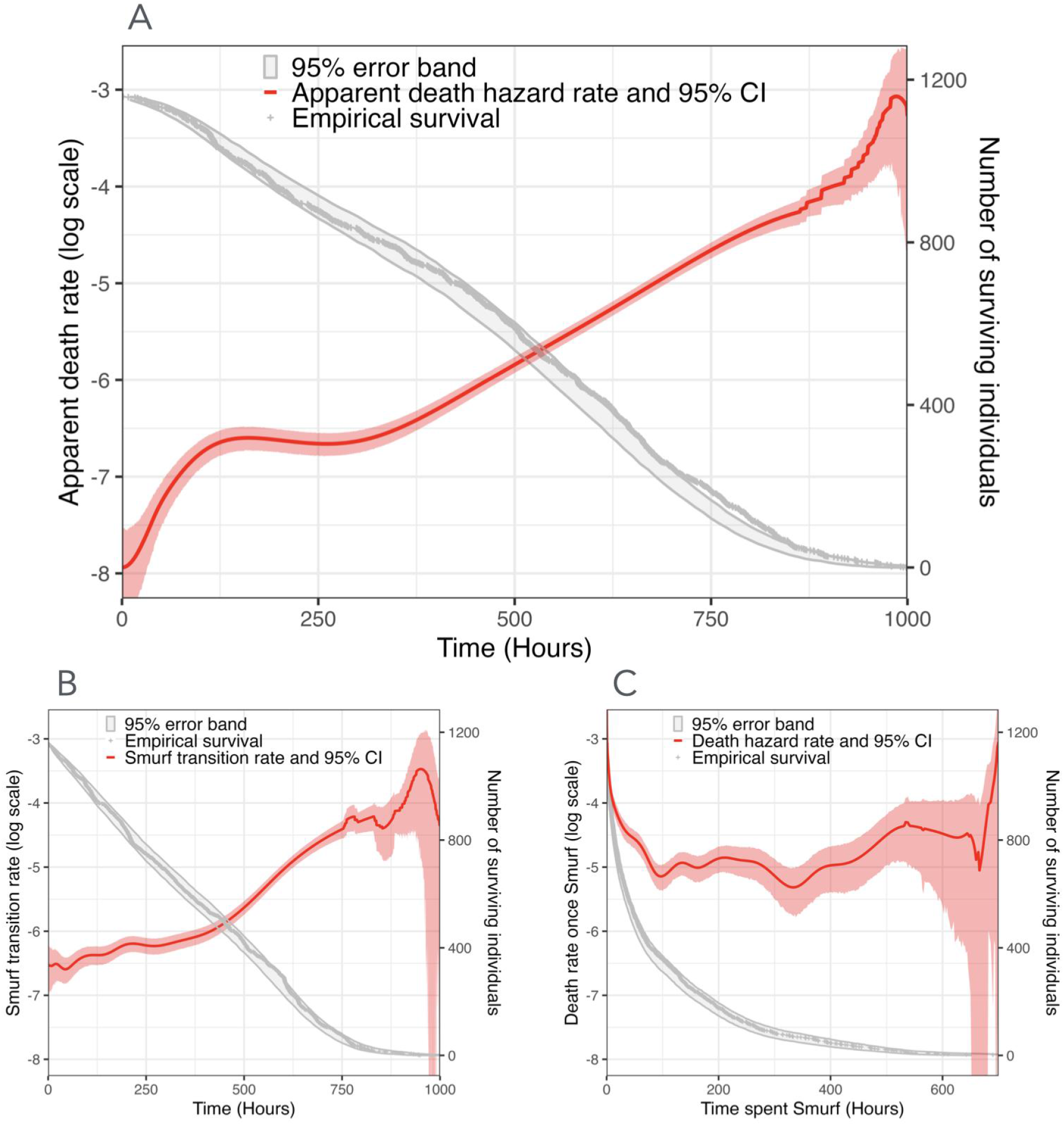
Non-parametric estimation of the apparent transition rates. A: Classic death rate estimate from birth to death. B-C: Two-phase transition rate estimates B: from non-Smurf to Smurf and C: from Smurf to death. In grey: empirical survival curves (grey dots) and 95% error band obtained by a bootstrap approach (light grey area). In red: transition rates (full line) and 95% asymptotic confidence intervals (light red area). A-B time 0 indicates day 11, age at transfer on the blue food.

The apparent death rate (Figure 2A), is increasing with the chronological age, which is not surprising as the process of ageing accelerates with age. We observe several successive contrasted tendencies: in log-scale, a fast increase followed by a plateau and then a linear increase - i.e., an exponential trend in linear scale. Many parameters are required to parametrise such a curve, whose mechanistic interpretation is difficult. The non-parametric estimation of the Smurf transition rate (Figure 2B) is also increasing with chronological age. This increase is slow before 400 hours and exponential subsequently.

Finally and more surprisingly, we observe in Figure 2B that once turned Smurf, flies first exhibit a high death rate, which then decreases. About 40% of flies die in the first 24 hours upon turning Smurf. The death rate decreases by about 78% during the first 24 hours, and then stabilizes at a high value before increasing again after 400 hours (around 16.5 days). It is unclear whether this final increase is biologically relevant or due to estimation uncertainty, as the confidence interval widens after 400 hours showing a lack of confidence in the results.

The time spent non-Smurf and the time spent Smurf have a slight negative linear correlation of -0.164 (see also Figure S3). To go further, Figure 3 shows the non-parametric estimation of the death rates when the flies are divided in four groups of equal sizes, based on the quartiles of the distribution of the time spent non-Smurf. Figure 3A shows a close-up for the first 50 hours spent Smurf, during which the death hazard rate is particularly high. The death rates in each subgroup have a similar shape. However, one can observe that the death rate of the flies which have spent the least time in the non-Smurf phase (their time spent non-Smurf is in the first quantile of the experimental distribution) is higher than the other quantiles between 5 and 24 hours, which is specifically the interval of Smurf age at which close to 40% of the flies die. However, such a higher death rate for a smaller time spent non-Smurf indicates a positive dependence, which comes in contradiction with the overall negative correlation coefficient observed.

**Figure 3.**
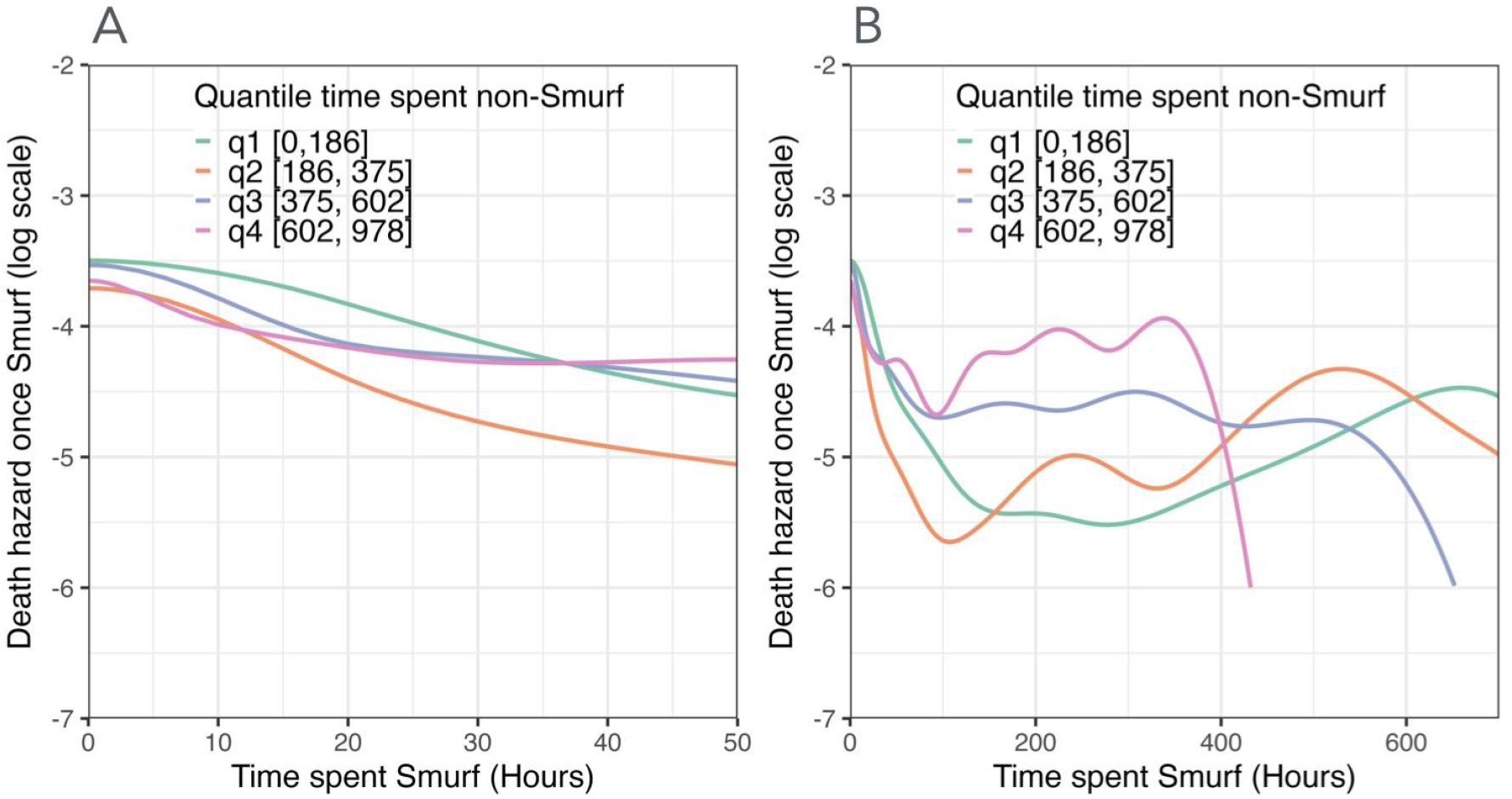
**Non-parametric estimation of the death rate once Smurf** with data separated according to the quartiles of time spent non-Smurf [0, 186], [186, 375], [375, 980]. A: Close-up between 0 and 50 hours. B: Full death rate between 0 and 750 hours.

### 2. Parametric modelling

Based on the observations detailed above, we propose a new model for the joint distribution of 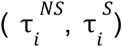, using the non-parametric estimations of Figure 2 to determine the most suitable parametric model. In the following, the chronological age is denoted by *t*, with *k*_*S*_ (*t*) the Smurf transition rate, and *a* the Smurf age, i.e. the time since the transition from non-Smurf to Smurf. To account for potential dependence between the durations of the two phases, we introduce the conditional death rate of Smurf flies *k*_*d*_ (*a, τ*^*NS*^), defined as the instantaneous rate at which a fly of Smurf age *a* dies, conditionally to the time *τ*^*NS*^ spent in the non-Smurf phase.

#### A Gompertz-Makeham law for the Smurf transition rate

As seen previously in Figure 2, the Smurf transition rate is relatively low - but nonzero - for small ages, and then increases. Among the four parametric models tested to describe the Smurf transition rate, the classical Gompertz-Makeham model^23^ *k*_*s*_ (*t*) = *f* + *g. exp*(*ht*) provided the best fit based on model selection criteria (see Supplementary Table S1 and Figure S1 for details on the comparison procedure). The three values of the parameters are fitted using maximum likelihood (see Supplementary methods). The fitted hazard rate and non-Smurf survival functions are shown on Figure 4, confirming the excellent fit of the Gompertz-Makeham model (see also Supplementary Quantile-Quantile (Q-Q) plots).

**Figure 4.**
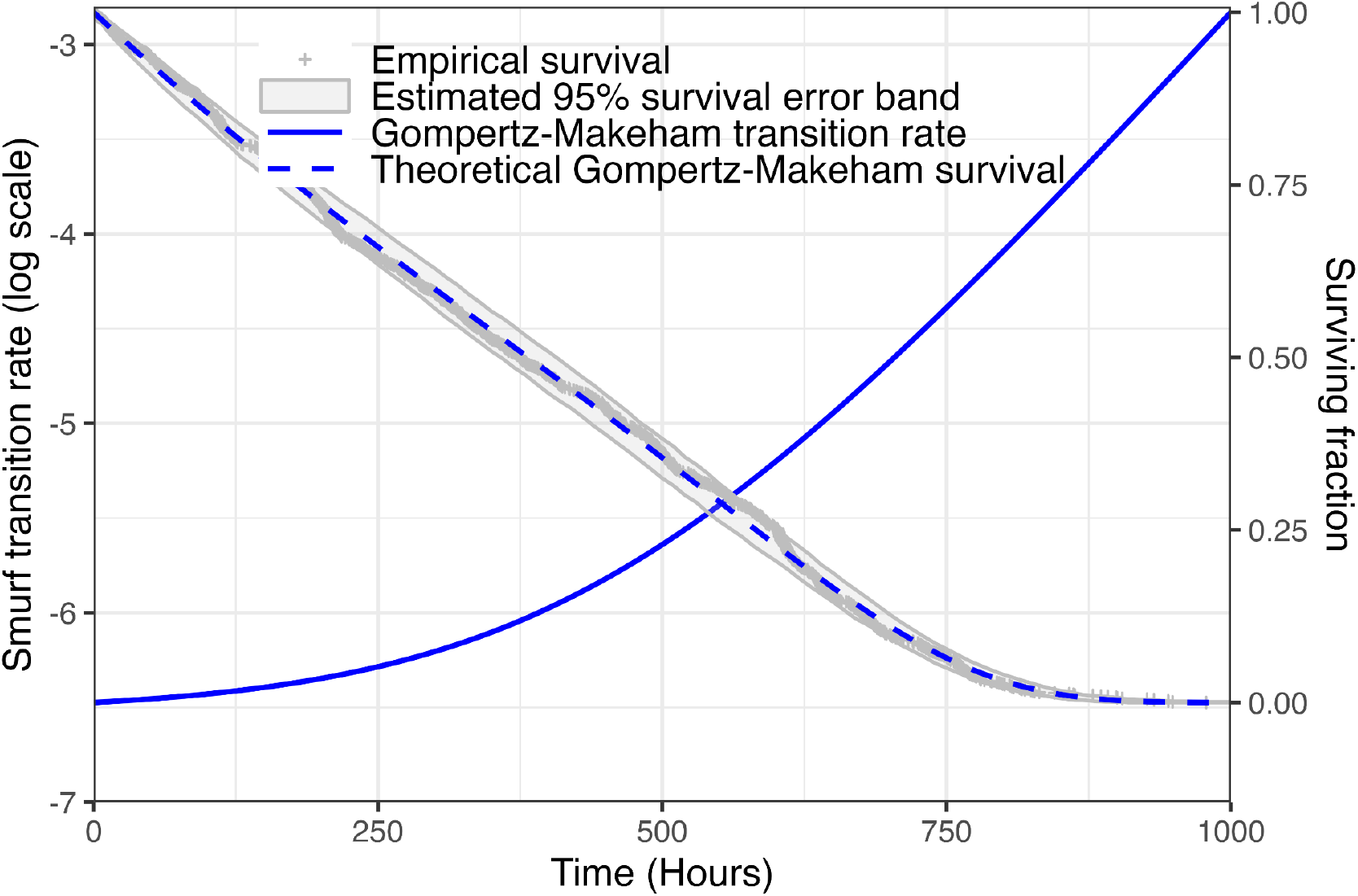
Best parametric choice for the Smurf transition rate. Blue line: best-fit transition rate using the Gompertz-Makeham parametrisation. Grey line: empirical survival curve. Light Grey area: 95% error band for the survival curve obtained by simulating with the best-fit transition rate (blue line). Time 0 indicates day 11, age at transfer on the blue food.

#### A decreasing death rate once Smurf, with dependence on the time spent non-Smurf

The full model for the joint distribution was selected from twelve candidate parametric models listed in Table 1. The conditional death rate once Smurf is parametrised by a Cox Model^24^, where the effect of the covariable (time spent as non-Smurf) is captured through an exponential coefficient. Formally, the death rate once Smurf can be written as *k*_*d*_ (*a, τ*^*NS*^) = *k*_*d*_ (*a*). *exp*(ɣ. *f*(*τ*^*NS*^)), where *a* denotes the Smurf age and *τ*^*NS*^ the time spent non-Smurf. The exponential coefficient captures the dependence, while the base hazard *k*_*d*_ (*a*) depends only on the Smurf age. We investigate three modelling scenarios: Models 1-3 assume no dependence between the time spent as non-Smurf and the time spent Smurf (ɣ =0), Models 4-6 incorporate dependence across all values of *τ*^*NS*^ and Models 7-12 introduce dependence only for *τ*^*NS*^> 200.

**Table 1.** Description of tested models. For a more detailed formulation see Table S4.

| Number | Dependence | $k_D$ | Number of parameters | BIC | Mean empirical Wasserstein distance |
| --- | --- | --- | --- | --- | --- |
| 1 | No | Constant | 4 | 12827 | 36.8 (2.02) |
| 2 | No | 2 Constants | 5 | 12577 | 29.7 (2.13) |
| 3 | No | Exponential | 6 | **12500** | 31.4 (2.25) |
| 4 | Yes | Constant | 5 | 12776 | 33.1 (2) |
| 5 | Yes | 2 Constants | 6 | 12551 | 26.4 (2.49) |
| 6 | Yes | Exponential | 7 | **12476** | 26.2 (2.22) |
| 7 | $\tau^{NS} > 200$ | Constant | 5 | 12752 | 30.9 (1.82) |
| 8 | $\tau^{NS} > 200$ | 2 Constants | 6 | 12535 | **24.5** (1.9) |
| 9 | $\tau^{NS} > 200$ | Exponential | 7 | **12463** | **24.4** (2.07) |
| 10 | $\tau^{NS} > 200$ | Constant separate fits | 7 | 12774 | 31.8 (2.07) |
| 11 | $\tau^{NS} > 200$ | 2 constants separate fits | 9 | 12516 | **23.00** (2.14) |
| 12 | $\tau^{NS} > 200$ | Exponential separate fits | 11 | **12431** | **23.02** (2.30) |

Within each model class, we test three parametrisations for the base hazard rate *k*_*d*_ (*a*) : constant, as hypothesised in Tricoire and Rera^10^ (Models 1, 4 and 7), piecewise constant Models 2, 5, 8 and 11), and a decreasing exponential form (Models 3, 6, 9 and 12), namely *k*_*d*_ (*a*) = *k*_1_ + *k*_2_ *exp*(− *d. a*). The latter functional shape matches that of the non-parametric estimations of Figures 2 and 3, and fits well the non-parametric estimator of the cumulated base hazard (see Figure S6, Model 6). Furthermore, regardless of the dependence structure or selection criterion (see Table 1, Figure S2, S4 and S5), the decreasing exponential form consistently outperformed the alternatives. This suggests either the existence of heterogeneity among Smurfs or that the higher mortality risk accompanying the transition is later coped with. Consequently, our analysis focuses on Models 3, 6, 9, and 12 to investigate the dependence between the time spent as non-Smurf and the time spent as Smurf.

#### Modelling the dependence

The negative correlation coefficient (-0.164) indicates a negative dependence between the times spent as non-Smurf and Smurf. The dependence is also visible on Figure 3, where it is shown that the death rate varies conditionally on the quantile of time spent non-Smurf. Not taking into account this dependence, as is done in Model 3, results in a poor fit of the Smurf population over time (see Figure S5) and the BIC and Wasserstein distance (Table 1).

A monotonous dependence pattern parametrised with the classical Cox model is introduced in Model 6. The Cox coefficient ɣ=7.33e-4 is estimated by maximum likelihood (see Supplementary for details), indicating a negative dependence between the durations of the two phases, consistent with the observed negative correlation: the longer the time spent as non-Smurf, the higher the death hazard rate and thus the shorter the time spent in the Smurf state. The Wald test assessing whether ɣ differs from zero yields a p-value of 5e-8, demonstrating that this dependence is statistically significant.

To go further, Table 2 reveals two distinct behavioural regimes in the joint distribution of time spent as non-Smurf and Smurf, depending on whether the time spent as non-Smurf falls below or above a threshold of 200 hours. This threshold was chosen as it is close to the first quartile of time spent as non-Smurf (186 hours) and corresponds to the value *x* at which the p-value of the Wald test reaches its minimum and the correlation coefficient is most negative, when restricting the analysis to the subset *τ*^*NS*^≥*x* (see Figure S4).

**Table 2.** **Statistical indicators of dependence between transition times** depending on the interval of time spent non smurf

| Time spent non-Smurf | Correlation coefficient | Mean time spent Smurf | Standard deviation of time spent Smurf | p-value of Wald test |
| --- | --- | --- | --- | --- |
| $\leq 200$ | -0.077 | 90.9 | 149.7 | 0.04 |
| $> 200$ | -0.28 | 97.6 | 112.1 | $6.3 \cdot 10^{-16}$ |

As shown in Table 3, flies that spent less than 200 hours as non-Smurf lived on average 91 hours Smurf, compared to 98 hours for those exceeding this threshold. In comparison, Model 6 predicts a greater expected time spent Smurf for flies that spent less than 200 hours non-Smurf. Moreover, the dependence between times spent in each phase spent non-Smurf is much weaker for individuals with *τ* ^*NS*^ ≤ 200, as indicated by both the Wald test, which is not significant and the correlation coefficient which is close to 0 (see Table 2). Overall, our estimate of the dependence is that the death rate increases by 3% for each day spent non-Smurf, confirmed by Raz et al. ^25^

**Table 3.** Statistical descriptors for experimental data vs selected Models 6 and 9. First line: mean (in hours), second line: standard deviation in parenthesis (in hours).

| Model | Time spent non-Smurf | Time spent Smurf | Time spent Smurf $\tau^{NS} \leq 200$ | Time spent Smurf $\tau^{NS} > 200$ | Life exp. $\tau^{NS} \leq 200$ | Life exp. $\tau^{NS} > 200$ | Life exp. |
| --- | --- | --- | --- | --- | --- | --- | --- |
| Data | 387.4<br>(237.3) | 92.8<br>(123.7) | 97.6<br>(49.7) | 91.0<br>(112.1) | 198.4<br>(156.6) | 589.1<br>(184.1) | 480.2<br>(248.9) |
| 6 | 386.9<br>(236.8) | 92.5<br>(125.6) | 134.1<br>(181.9) | 83.1<br>(111.5) | 215.8<br>(160.3) | 580.5<br>(197.8) | 479.3<br>(249.2) |
| 9 | 386.9<br>(236.8) | 90.9<br>(127.2) | 88.7<br>(148.1) | 92.7<br>(128.1) | 185.7<br>(131.1) | 633.9<br>(39.3) | 509.5<br>(214.9) |

Based on these observations, we introduce Model 9 in which no dependence is assumed for individuals with *τ* ^*NS*^ ≤ 200, (i.e. a Cox coefficient equal to 0 on this interval), while a non-zero Cox coefficient is fitted using only the subset of individuals with *τ*^*NS*^ > 200 (see Table 1 and Table S4), hence *k*_*d*_ (*a, τ*^*NS*^) = *k*_*d*_ (*a*). *exp*(ɣ. *f*(*τ*^*NS*^) ), with *k*_*d*_ (*a*) a decreasing exponential function and 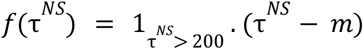-where *m* is a centering constant (typically, the empirical mean of the covariable). In the Supplementary material, we present a further extension where the piece-wise dependence of 9 is retained, but the base hazard rates are fitted separately on these two intervals (see Table 1 and Table S4, Model 12).

The means and variances of lifetimes and time spent in the non-Smurf and Smurf phases are reported in Table 3 for Models 6 (uniform dependence) and 9 (piece-wise dependence). According to the table, Model 6 provides a better estimate of the mean life time, but lacks precision in rendering the laws of time spent Smurf and time spent non-Smurf for time spent non-Smurf under 200 hours, in comparison with Model 9. The Smurf population dynamics predicted by Model 9 also lead to a better fit to the experimental data, as shown in Figure 5. Model 12 lacks the generality offered by Model 6 and to some extent Model 9, and carries a higher risk of overfitting, as it may capture dataset-specific features rather than robust biological mechanisms.

**Figure 5.**
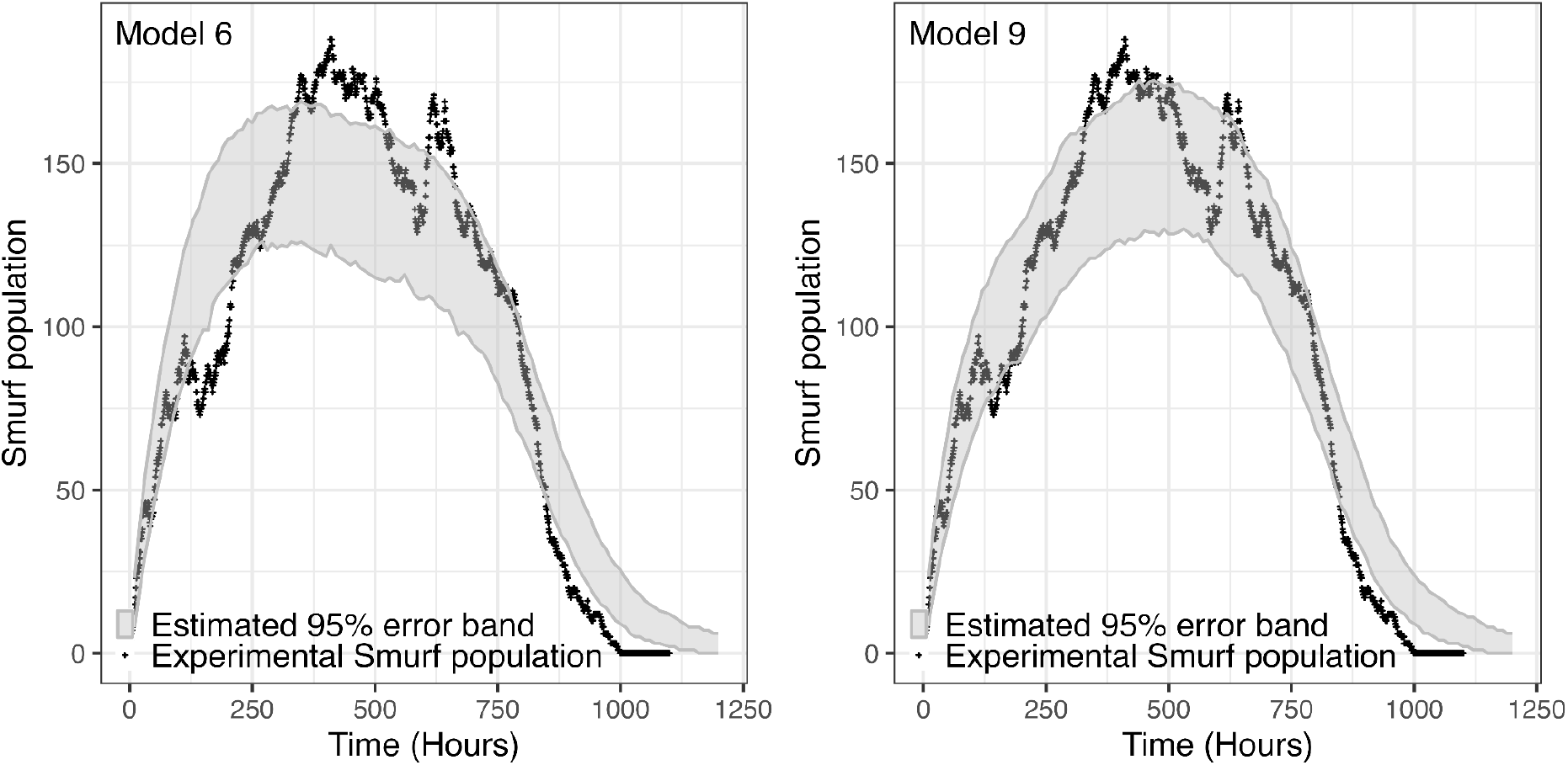
Experimental vs modelled Smurf population over time. Taking into account the higher mortality of early (<200h) Smurfs allows a better fit of experimental data with our model. Dots: experimental data. Grey surface: 95% estimated error bands for Models 6 (left) and 9 (right). Time 0 indicates day 11, age at transfer on the blue food.

Finally, the death rates once Smurf for different durations spent non-Smurf are presented in Figure 6. The graph on the left showcases that introducing dependence through the Cox model in Model 6 produces a vertical shift of the death rate curves: the longer a fly remains in the non-Smurf state, the higher its death rate once Smurf. Consequently, flies that spend more time non-Smurf are predicted to spend less time in the Smurf state on average. Conversely, Model 9 predicts a higher death rate for individuals with a shorter time spent as non-Smurf, which may reflect an underlying frailty in these individuals.

**Figure 6.**
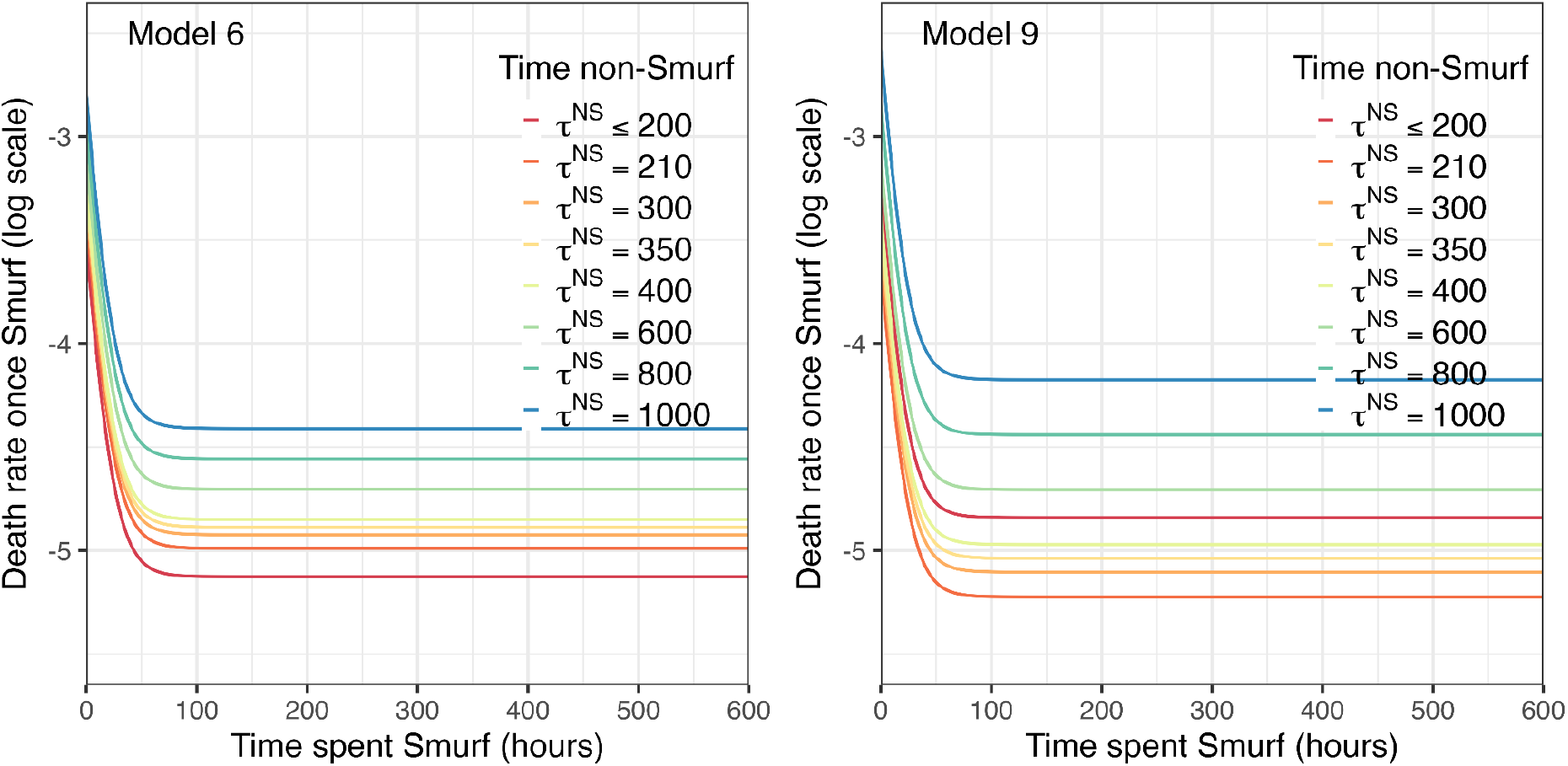
**Death rate once Smurf for different times spent non-Smurf** in Models 6 (left) and 9 (right).

### 4. Extension and validation of the model

#### Comparison with one-phase models: prediction of lifetime distribution

Up to this point, our study has focused on modelling the joint distribution of time spent non-Smurf and time spent Smurf, by analysing both hazard rate separately and by estimating the dependence. While this approach yields detailed models that accurately reproduce the marginal distributions of time spent in each phase, these models have not yet been evaluated in terms of their ability to predict total lifetime.

The density histogram of total experimental lifetimes presented in Figure 7 A reveals a bimodal distribution, with a first mode around 125 hours and a second, more prominent mode around 600 hours, close to the expected lifespan of the fly strain used in this study. As presented in Figure 2A, this leads to an apparent death rate with a bump around 125 hours, followed by a plateau. To assess the relevance of the two-phase framework, we compare Models 6 and 9 with one-phase models with parametrisations of mortality rates commonly used in survival analysis: Gompertz-Makeham^23^, Gamma-Gompertz^26^ and Generalised Gamma^27^ . To quantify the fits, we present 6 indicators: the visual agreement of the lifetime density (Figure 7 A) and hazard rate (Figure 8 B), Q-Q plots (Figure S7) and three quantitative measures, the BIC, 1D Wasserstein distance and log-likelihood (see Table S2).

**Figure 7.**
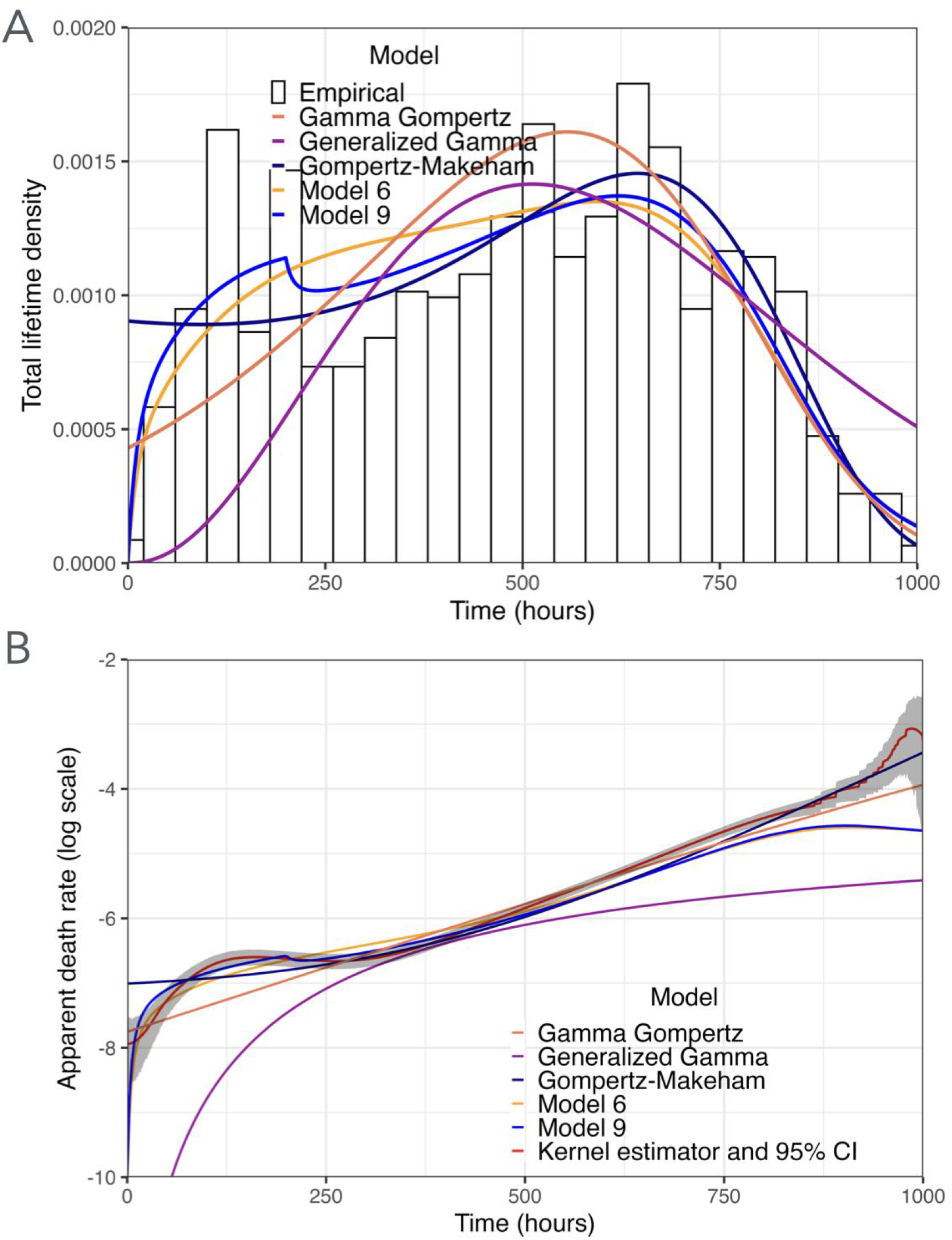
Experimental versus modelled lifetimes. (A) Histogram of densities for Models 6, 9 and three models of one-phase death hazard rate: Gompertz-Makeham, Gamma-Gompertz and Generalised Gamma. (B) Apparent death rate. Comparison of non-parametric estimation with modelled death rates for Models 6, 9, and three models of one-phase death hazard rate: Gompertz-Makeham, Gamma-Gompertz and Generalised Gamma. Time 0 indicates day 11, age at transfer on the blue food.

**Figure 8.**
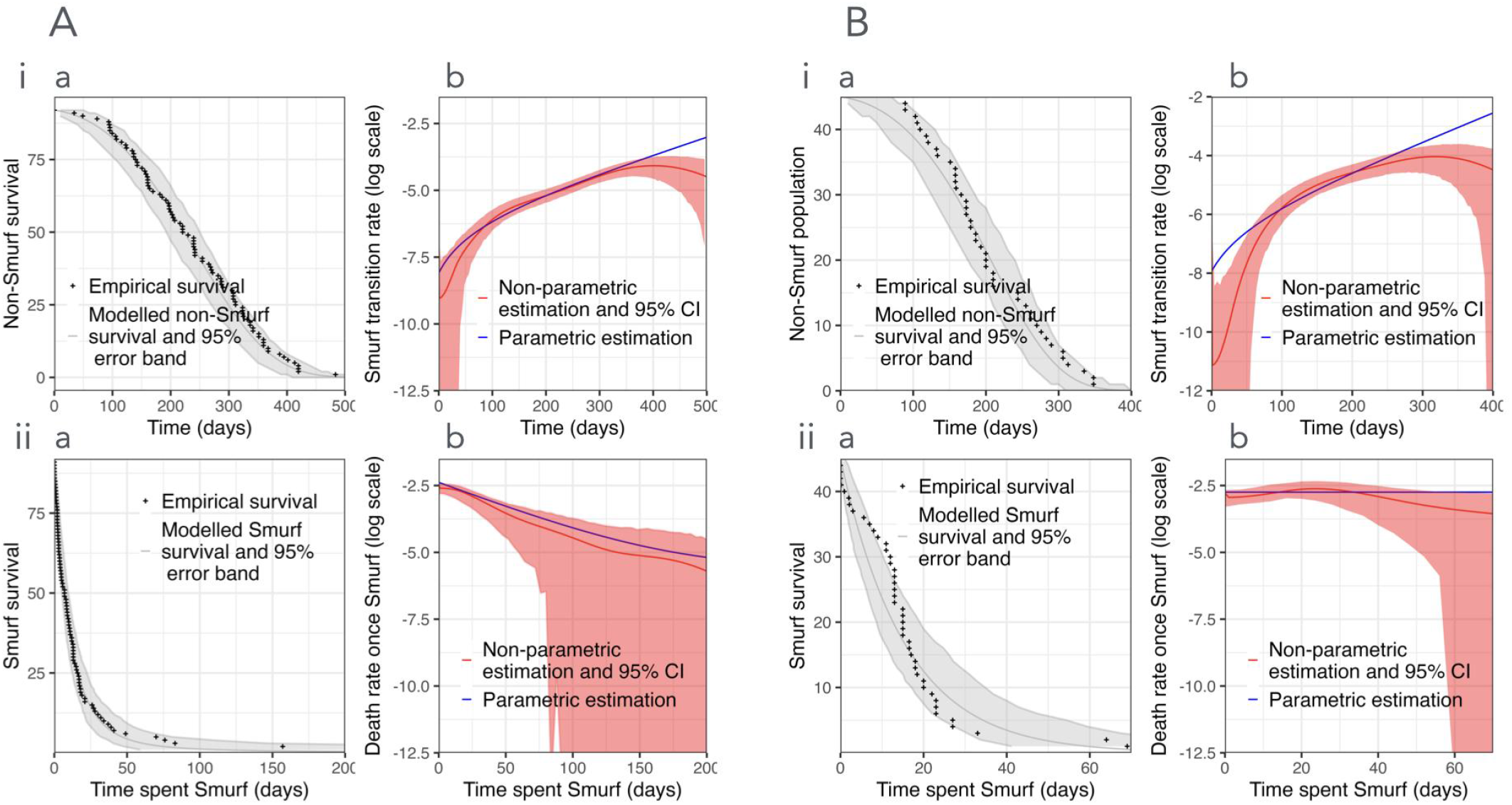
**Fly-based two-phase model applies to mice Smurf survival analysis** for two laboratory strains, C57BL6/J (A) and AKR/J (B). Non-parametric (red) and parametric (blue) estimations of non-Smurfs (i) and Smurfs (ii) survival functions (a) and hazard rates (b). Panels i, time 0 indicates the age when we started monitoring individuals, 94 days for AKR/J and 661 days for C57BL6/J.

Figure 7 A presents the modelled densities of lifetimes for Models 6 and 9 along with the three parametrisations of a one-phase mortality rate. It is apparent that Model 9 is the only one which renders the bimodality of lifetimes, even though it was not explicitly designed to do so. Notably, it achieves this with relatively few components and standard hazard rate formulations: a Gompertz-Makeham hazard for the non-Smurf to Smurf transition, a decreasing exponential baseline hazard, and a Cox-type dependence for the death rate after Smurf transition. By contrast, reproducing bimodality within a single-phase framework would require substantially more complex hazard functions, thereby reducing the scope for mechanistic interpretation. Although it does not present bimodality, Model 6 also renders a high density for the first 250 hours, in line with the experimentally high apparent death rate.

Both the BIC and the Wasserstein distance indicate that the Gompertz-Makeham parametrisation of the apparent death rate is the closest to the data (see Table S2) While the log-likelihood favors Model 12, this criterion does not account for overfitting. In comparison, the penalisation introduced by the BIC results in a poorer performance of Model 12, thus indicating a potential overfitting. Among the three one-phase mortality-rate parameterizations considered, Gompertz–Makeham clearly performs best for our dataset. This could be explained by the fact that the Smurf phase is relatively short compared to total lifespan, and the one-phase Gompertz-Makeham parametrisation primarily captures the dynamics of the non-Smurf phase.

According to the Q-Q plots (Figure S7), Model 9 is a good overall fit, but lacks precision beyond 800 hours in comparison with the one-phase Gompertz-Makeham model. However, the behavior of the distribution tails was not a primary focus of this study. In particular, Figure 2 suggests an increase in the death rate of Smurf flies after approximately 700 hours, a feature that is biologically plausible in survival analysis but would require more complex parameterisations to capture within our modelling framework. Given the limited reliability of the data at such advanced ages, we choose to focus on the bulk of the lifetime distribution. As a consequence, our models tend to overestimate the lifetimes of the oldest flies, as visible in Figure S7.

The two-phase models present a notably better fit for the early deaths, when the lifetime density rises steeply. Introducing two phases allows the density of lifetimes to increase rapidly from zero: initially, all flies are in the non-Smurf state and no death occurs, and deaths begin only once Smurf flies appear. In contrast, one-phase Gompertz–Makeham models assume a direct transition from life to death, so reproducing early deaths requires a high mortality density from time zero.

At this stage, it remains unclear whether this apparent bimodality, characterised by a significant number of flies dying before 250 hours is biologically meaningful, or simply specificities of our dataset. Potential outside causes for this bimodality are discussed in Section Discussion.

#### Generalisation to mice survival using Smurf data

To further validate our model and our approach, we test it on a dataset of mice survival, taken from Cansell et al.^15^ As what was done for the Drosophila data, non-parametric estimations allow to get a first unbiased idea of the rates, while parametric estimations provide a more quantitative and comparable description of its dynamics. Unlike in Drosophila, Smurf status in mice cannot be assessed directly through visual detection of a blue food dye. Instead, individuals are monitored for changes in several indicators, including inflammation, metabolic parameters, microbiota composition, and intestinal permeability. The age at which these indicators shift is used to determine the transition to the Smurf state. The processing of the data is detailed in the Supplementary methods. The Smurf assays were performed for two different mice strains with different life expectancies, for which we fit the parameters independently. After treatment, the dataset for the AKR/J mice strain is of size 45 and for the C57BL6/J mice of size 92. Due to the low sample sizes, we fit Model 3 only, where no dependence is assumed between the times spent in both phases and the death rate of Smurf individuals is modeled with a decreasing exponential.

The results of the non-parametric and parametric estimations of the Smurf transition rate and death rates are presented in Figure 8, along with confidence intervals and survival functions, to assess goodness of fit. Figure 8A and 8B both show that the Gompertz-Makeham parametrisation for the Smurf transition rate yields a good fit, most particularly for the C57BL6/J mice in Figure 8A. As for the death rate once Smurf, the decreasing exponential rate seems relevant for C57BL6/J mice (A), but results in a constant rate for AKR/J mice with the coefficient of the exponential being fit to nearly 0. Qualitatively, the non-parametric estimator of the death rate for AKR/J mice (B) seems to be increasing and then decreasing, which is a behaviour that could not be rendered with a decreasing exponential parametrisation. However, the constant death rate is within the confidence interval of the non-parametric estimation for the first 50 days, which captures the bulk of the data as only 2 mice die after 50 days spent Smurf. Overall, the qualitative behaviour of the Smurf transition rate is consistent with our conclusions on Drosophila, and the death rate for C57BL6/J mice also presents the same shape as for drosophila. The fit is poorer for AKR/J mice, but it is hard to interpret due to the small sample size.

## Discussion

In this article, we conducted a thorough statistical analysis of the data published in Tricoire and Rera^10^, based on a rigorous statistical framework to test and refine the two-phase ageing model. Building on insights from non-parametric estimation, we developed a parametric model that reveals three principal findings that complement and extend previous results. These findings provide key ingredients that a mechanistic model should incorporate.

### The Smurf transition follows a Gompertz-Makeham law

The transition rate from non-Smurf to Smurf was previously approximated using a linear equation^10^. We show that a Gompertz-Makeham parametrisation provides a significantly better fit to the data. The transition to Smurf is not a completely stochastic failure, i.e. independent of age, as it reflects progressive underlying deterioration with the risk of transitioning to intestinal barrier dysfunction increasing exponentially with age. The discontinuous two-phase model explains classical continuous conceptions of ageing: the phases are discrete, but the population-based apparent exponential mortality increase results from a combination of the rate of transitioning between phases and the subsequent high mortality rate of Smurfs.

Importantly, since the Smurf phase is relatively short compared to the total lifespan, it minimally impacts classical survival curves. The Gompertz-Makeham parametrisation that best describes total lifetime in single-phase models primarily captures the dynamics of the non-Smurf phase, where the continuous component of ageing resides.

### Acute vulnerability upon Smurf transition

Our most striking finding is the remarkably high mortality immediately following the Smurf transition: approximately 40% of flies die within 24 hours, after which the death rate declines before stabilizing towards a constant value. This pattern was overlooked under the constant-rate assumption^10^ and refines the predictability of death for Smurf individuals. The constant death rate previously described remains valid for “sufficiently old” Smurfs, see Figure 6. Our modification concerns the acute phase immediately post-transition. Interestingly, both phenomena are captured by the mechanistic saturating removal model proposed by Raz et al^25^ .

We interpret this finding as evidence that the physiological reorganisation accompanying the Smurf transition carries its own mortality risk, a moment of systemic vulnerability distinct from the pre-existing decline that triggered the transition. A fly surviving the first 24 hours as Smurf has substantially better short-term prognosis than one that just transitioned. Our 24-hour observation interval may actually underestimate the concentration of this mortality peak, and higher temporal resolution could characterise the acute vulnerability phase more precisely. In addition, the suggested increase in death rate after 700 hours spent Smurf (Figure 2C) might reflect genuine late-life acceleration of mortality or solely estimation uncertainty as sample size diminishes.

### Dependence between phases

The weak negative correlation between time spent non-Smurf and subsequent Smurf survival (r = -0.164), captured through a Cox model, indicates that flies transitioning late die faster once Smurf. This is consistent with continuous ageing operating within the non-Smurf phase: a fly remaining non-Smurf for 600 hours has accumulated more sub-threshold damage than one transitioning at 200 hours, even though both are phenotypically indistinguishable. This aligns with transcriptomic evidence that old non-Smurfs have gene expression profiles approaching those of Smurfs^7^ and the mechanistic modeling proposed by Raz et al^25^.

The absence of this dependence among early-transitioning flies ( *τ*^*NS*^ < 200 hours), combined with their higher early mortality, suggests further biological heterogeneity among Smurfs. These individuals may transition through mechanisms distinct from progressive ageing, i.e. developmental defects, acute injuries, or stochastic barrier failures, and their Smurf-phase mortality is dominated by whatever caused the premature transition rather than accumulated ageing damage. It was indeed previously shown that lifespan-reducing mutations would trigger early mortality through Smurf transition^6^.

### Emergent bimodality from simple components

Model 9 produces bimodal lifetime distributions without being explicitly designed for this purpose. The early mode around 125 hours and the later mode around 600 hours emerge from the biologically interpretable piecewise Cox dependence. Rendering such bimodality with a single-phase model would require hazard functions of complex parametric form, without straightforward interpretation. However, it is not clear whether this bimodality is robust across genetic backgrounds or specific to our dataset as it could reflect experimental perturbations rather than intrinsic ageing dynamics. Experimental causes could also explain the absence of dependence observed on early-transitioning flies (*τ*^*NS*^ ≤ 200 hours).

### Application to other survival data

Our analysis derives from a single genetic background with individual observations, and extension to other strains, males, or natural populations requires further investigation. As a first validation, we present a fit of the 2-phase model on DGRP line 377 in Figure S8. For line 377, only population-level data are available: the time spent in each phase is unknown, with only total lifetimes and a few observations of Smurf proportions recorded. Inferring model parameters from incomplete population data is mathematically challenging. In order to overcome this difficulty, we used the parametric Model 6 which can be fitted using this population data by fitting only the Smurf transition rate. This approach results in a very good fit of the population survival and Smurf proportion. This shows the robustness of our study, but would benefit from more complete and precise data on different genetic backgrounds to assess to what extent the conclusions of our study can be generalised, particularly concerning the death hazard rate once Smurf. Fitting a 2-phase model using population data rather than individual data would require further theoretical investigations from a statistical perspective. Multi-strain analysis would allow a better assessment of the dependence structure. Intervention studies could deploy this framework to decompose treatment effects: does a lifespan-extending manipulation reduce the Smurf transition rate *k*_*S*_, the death rate *k*_*d*_ or the dependence parameter *γ*? Each would suggest different mechanisms and possibly distinct genes involved.

We also applied the model to the mouse Smurf data we recently published^15^, to provide preliminary cross-species validation. We used the same methodology on this second dataset, with a complementary use of non-parametric and parametric estimations to best describe the data. The transposition of the 2-phase model initially calibrated on Drosophila data to mouse data shows that they follow a similar survival dynamics: an initial phase during which there is an increasing probability with chronological age of entering the Smurf phase followed by the Smurf phase, characterized by systemic vulnerability with little dependence on the chronological age. The fit is less accurate for AKR/J mice, though difficult to interpret given the small sample size. An important limitation is that mouse Smurf status must be inferred from composite biomarkers rather than direct dye visualisation, introducing a measurement uncertainty not present in the Drosophila dataset.

### Toward a broader framework: discontinuous ageing in humans?

In demographic modelling, population heterogeneity is often captured through the notion of frailty. The Smurf phenotype offers a measurable alternative as heterogeneity is directly observable rather than inferred. It is important to note that the dependence structure we find, although weak, suggests residual heterogeneity within phases.

Recent multi-omics data suggest that discontinuous ageing dynamics may extend to humans. Shen and colleagues^28^ performed comprehensive profiling on a longitudinal cohort of 108 participants aged 25-75 years and found consistent nonlinear patterns in molecular markers, with substantial dysregulation occurring at approximately 44 and 60 years of chronological age. These two waves of molecular changes were robust across different statistical cutoffs, highlighting their biological significance. Distinct functional pathways were associated with each period: immune regulation and carbohydrate metabolism shifted during the 60-year transition, while cardiovascular disease markers and lipid and alcohol metabolism changed at the 44-year transition^28^.

These findings resonate with our two-phase model, though the relationship is not straightforward. The Smurf transition in flies represents a single discrete event preceding death, whereas the human data suggest multiple waves of molecular reorganisation across the lifespan. One interpretation is that humans experience several transitions analogous to the Smurf phenotype, each marking entry into a new physiological regime. Alternatively, the 44-,60- and 80-year ageing waves may reflect population-level averages of heterogeneous individuals. These time points would correspond to populations with different proportions of “Smurf” individuals, increasing with chronological age.

A limitation of the human multi-omics study is its cross-sectional design within age bands: comparisons across individuals of different ages may capture some biological ageing effects, but longitudinal tracking of the same participants would be needed to confidently identify discrete transitions within individuals. Nevertheless, the convergence of evidence - nonlinear molecular dynamics in humans, discrete phenotypic transitions in model organisms - strengthens the case that ageing proceeds through qualitatively distinct phases rather than continuous decline. A phenomenon that seems to be detected through methods independent from ours in mice^29^ and Drosophila^30^.

### Connection to mathematical models of damage accumulation

This study relies on recent non-parametric statistical work^22^, allowing us to capture precisely the value of a hazard rate near t=0, when the hazard rate is different from 0. This methodology allowed us to identify the high death rate once Smurf, which could not have been detected with classical methods due to their well-known boundary bias. This feature should be reproduced by a relevant mechanistic model of the Smurf phenotype, as in Raz et al^25^. Our statistical framework is also coherent with recent theoretical work on ageing dynamics. Yang and colleagues^31^ developed, independently from us, what they termed the saturated removal (SR) model, a stochastic differential equation for damage production and removal, calibrated using single-cell damage measurements in *E. coli*. In this model, noise is amplified by a rising production of damage that saturates its own removal mechanisms^32^.

A striking prediction of the SR model is “shortening twilight”: the remaining lifespan after a given damage threshold is crossed decreases with the age at which that threshold is reached. This maps directly onto our finding of (slight) negative dependence between time spent non-Smurf and subsequent Smurf survival. The SR model also offers a framework for understanding intervention effects. Our two-phase framework could help operationalise these predictions. Assessing which of the Smurf transition rate *k*_*S*_, the Smurf death rate *k*_*d*_, or the dependence parameter γ is affected by an intervention would bring different implications for healthspan compression.

Interestingly, Alon’s group found that relative variation in damage among *E. coli* cells drops with age - cells become more similar in terms of damage, indicating increasing determinism as death approaches. This parallels our observation that early-transitioning flies (*τ*^*NS*^≤ 200 hours) show high variability and no dependence structure, while late-transitioning flies exhibit tighter coupling between phases. Early transitions may be dominated by stochastic events or individual-specific defects. Late transitions may reflect the increasingly deterministic dynamics of saturated repair. It is however important to keep in mind here that the coupling remains moderate and the remaining life expectancy of Smurfs is mostly unaffected by the age at transition.

### Conclusion and Perspectives

The convergence between our empirical findings in flies, the multi-omics waves in humans, and the mathematical framework of saturated damage removal suggests a potential universality in ageing mechanisms. The Smurf phenotype may represent a visible manifestation of the transition from a regime where repair can keep pace with damage to one where repair is overwhelmed, what Clarke and Smith^4^ intuited in 1961 as the division between “ageing” and “dying”. In addition, the broad evolutionary conservation of this discontinuous process suggests that it is an early invention of living organisms, capable of reproduction and homeostasis^33,34^.

The statistical methodology introduced in this paper can be applied to other datasets and, more generally, to the estimation of transition rates in multi-phase ageing models, with the advantage of providing point estimates and confidence intervals, critical when dealing with small sample sizes or mild effects. Future work should test whether the two-phase model displays similar dependence in other organisms, and whether the molecular signatures of the Smurf transition overlap with the waves of ageing identified in humans. Raz et al^25^ shows that such discontinuous models describe well individual healthspan trajectories in humans using frailty index as a surrogate for Smurfness. The intestinal permeability biomarkers already validated in clinical settings^20,35^ offer a path toward validating Smurfness in frail human populations. Using the methodology introduced in this paper, a similar systematic approach could also be applied to existing human data on pathologies or multidimensional frailty measures, in order to identify discontinuities in ageing.

## Methods

All of the simulations are run by using the R package IBMPopSim^36^ and can be found at the following link https://github.com/luce-breuil/smurf_stat

- **Statistical inference :** The non-parametric estimations were done following the theoretical results and the methodology developed in Breuil and Kaakaï^22^. The parametric estimations were done with maximum likelihood estimation as detailed in the Supplementary methods. The indicators we use to compare the models are as follows:
- **The Bayesian Information criterion (BIC)**. For a parametrization *k* with parameters θ, the BIC is defined by 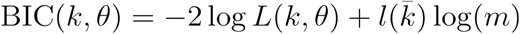 where *l* is the number of fitted parameters, which depends on the chosen parametrization, and *m* the sample size. *L*(*k*, θ) is the log-likelihood of the model with hazard rate *k* and parameters θ. This criterion penalizes models that have a lot of parameters to prevent over-fitting^37^ .
- **The empirical Wasserstein distance** between the distribution of the data and the model distribution. To compute the Wasserstein distances, we use the functions wasserstein and wasserstein1D of the package transport in R^38^. We consider the mean empirical distance between the data and 100 simulations of a given model. See Supplementary methods for details.
- **Q-Q plots**, or quantile-quantile plots, computed on a simulated population of 12000 individuals.
- **95% error bands/Confidence intervals**. The error bands shown in several of our figures are done by simulating 500 populations and plotting pointwise the 2.5% and 97.5% quantiles of the quantity of interest over the 500 simulations. The confidence intervals shown (for parametric and non-parametric estimations), are computed using their asymptotic theoretical expressions (see Supplementary methods). As the number of individuals decays with time, the confidence intervals widen, showing the lack of confidence we expect when only few individuals remain alive at the time of estimation.
- **Visual comparison of survival function**. The use of the non-Smurf survival function as an indicator of goodness-of-fit is done since it is a quantity that can be represented empirically from the data without preprocessing, contrarily to what would be necessary for the density and the transition rate.

## Funding and acknowledgments

M. Rera is funded by ANR-20-CE44-0010 and the authors by the ANR project ANR-25-CE45-5355. The authors would like to thank the Chaire Modélisation Mathématique and Biodiversité Veolia Environnement - Ecole Polytechnique - Muséum National d’Histoire Naturelle - Fondation X for fostering this collaboration and partial funding. The research of Sarah Kaakai was funded by the European Union (ERC, SINGER, 101054787). Views and opinions expressed are however those of the author(s) only and do not necessarily reflect those of the European Union or the European Research Council. Neither the European Union nor the granting authority can be held responsible for them.

