## Supplementary Figures for "Acute Smurf mortality and age-dependence in a two-phase ageing model: Statistical inference from drosophila and mice experimental data"

### Supplementary tables

| $k_S$ | Parameters | BIC | Wasserstein distance (SD) |
|---|---|---|---|
| Polynomial | 2 | 15819 | $46.5\,(5.6 \cdot 10^{-3})$ |
| Gompertz | 2 | 15729 | $23.2\,(3.0 \cdot 10^{-3})$ |
| Gompertz-Makeham | 3 | **15683** | $\mathbf{10.3\,(2.1 \cdot 10^{-3})}$ |
| Weibull | 2 | 15917 | $43.8\,(3.3 \cdot 10^{-3})$ |

Table S1: **Comparison of parametric models for the Smurf transition rate** with the BIC and 1D Wasserstein distance: polynomial, Gompertz, Gompertz-Makeham and Weibull. The minimum value is in bold font.

| Model | Number of parameters | BIC | log-likelihood | Wasserstein distance (SD) |
|---|---|---|---|---|
| 2-phase model 6 | 7 | 15941 | $-7946$ | 20.5(4.0) |
| 2-phase model 9 | 8 | 15926 | $-7935$ | 15.7(3.5) |
| 2-phase model 12 | 11 | 15936 | $\mathbf{-7929}$ | 12.9(3.3) |
| Gompertz-Makeham | 3 | **15890** | $-7935$ | **12.2(2.8)** |
| Gamma-Gompertz | 3 | 15941 | $-7960$ | 25.4(3.7) |
| Generalized gamma | 3 | 16163 | $-8071$ | 153.1(9.5) |

Table S2: **Comparison of parametric models of one-phase death hazard rate** with BIC, 1D Wasserstein distance and log-likelihood.

| Model | Expression | Parameter | Parameter value |
|---|---|---|---|
| Polynomial | $k_S(t) = a + b \cdot t$ | $a$ | $6.0 \cdot 10^{-9}$ |
| | | $b$ | $1.9 \cdot 10^{-3}$ |
| Gompertz | $k_S(t) = ce^{dt}$ | $c$ | $1.0 \cdot 10^{-3}$ |
| | | $d$ | $2.9 \cdot 10^{-3}$ |
| Weibull | $k_S(t) = \frac{v}{w}\left(\frac{t}{w}\right)^{v-1}$ | $v$ | 1.6 |
| | | $w$ | 435 |
| Gompertz-Makeham | $k_S(t) = f + ge^{ht}$ | $f$ | $1.5 \cdot 10^{-3}$ |
| | | $g$ | $7.5 \cdot 10^{-5}$ |
| | | $h$ | $6.6 \cdot 10^{-3}$ |

Table S3: **Parameter values for parametric models of Smurf transition rate.**

| Model | Expression | Parameter | value |
|---|---|---|---|
| Model 1 | $k_D(t) = k$ | $k$ | $1.01 \cdot 10^{-2}$ |
| Model 2 | $k_D(t) = k_1 \mathbb{1}_{t\leq 24} + k_2 \mathbb{1}_{t>24}$ | $k_1$ | $2.2 \cdot 10^{-2}$ |
| | | $k_2$ | $8.0 \cdot 10^{-3}$ |
| Model 3 | $k_D(t) = k_1 + k_2 e^{-dt}$ | $k_1$ | $7.3 \cdot 10^{-3}$ |
| | | $k_2$ | $2.9 \cdot 10^{-2}$ |
| | | $d$ | $6.7 \cdot 10^{-2}$ |
| Model 4 | $k_D(t,u) = k e^{\gamma(u-\mu)}$ | $k$ | $1.1 \cdot 10^{-2}$ |
| | | $\gamma$ | $7.3 \cdot 10^{-4}$ |
| | | $\mu$ | 387 |
| Model 5 | $k_D(t,u) = (k_1 \mathbb{1}_{t\leq 24} + k_2 \mathbb{1}_{t>24}) e^{\gamma(u-\mu)}$ | $k_1$ | $2.2 \cdot 10^{-2}$ |
| | | $k_2$ | $8.3 \cdot 10^{-3}$ |
| | | $\gamma$ | $7.33 \cdot 10^{-4}$ |
| | | $\mu$ | 387 |
| Model 6 | $k_D(t,u) = (k_1 + k_2 e^{-dt}) e^{\gamma(u-\mu)}$ | $k_1$ | $7.8 \cdot 10^{-3}$ |
| | | $k_2$ | $3.1 \cdot 10^{-2}$ |
| | | $d$ | $7.9 \cdot 10^{-2}$ |
| | | $\gamma$ | $7.33 \cdot 10^{-4}$ |
| | | $\mu$ | 387 |
| Model 7 | $k_D(t,u) = k e^{\gamma(u-\mu)\mathbb{1}_{u\geq 200}}$ | $k$ | $1.1 \cdot 10^{-2}$ |
| | | $\gamma$ | $1.3 \cdot 10^{3}$ |
| | | $\mu$ | 498 |
| Model 8 | $k_D(t,u) = (k_1 \mathbb{1}_{t\leq 24} + k_2 \mathbb{1}_{t>24}) e^{\gamma(u-\mu)\mathbb{1}_{u\geq 200}}$ | $k_1$ | $2.2 \cdot 10^{-1}$ |
| | | $k_2$ | $8.5 \cdot 10^{-3}$ |
| | | $\gamma$ | $1.3 \cdot 10^{3}$ |
| | | $\mu$ | 498 |
| Model 9 | $k_D(t,u) = (k_1 + k_2 e^{-dt}) e^{\gamma(u-\mu)\mathbb{1}_{u\geq 200}}$ | $k_1$ | $7.9 \cdot 10^{3}$ |
| | | $k_2$ | $3.1 \cdot 10^{-2}$ |
| | | $d$ | $8.0 \cdot 10^{-2}$ |
| | | $\gamma$ | $1.3 \cdot 10^{3}$ |
| | | $\mu$ | 498 |
| Model 10 | $k_D(t,u) = \mathbb{1}_{u\geq 200} k_1 e^{\gamma(u-\mu)} + k_2 \mathbb{1}_{u<200}$ | $k_1$ | $1.0 \cdot 10^{-2}$ |
| | | $k_2$ | $1.1 \cdot 10^{-2}$ |
| | | $\gamma$ | $1.3 \cdot 10^{3}$ |
| | | $\mu$ | 498 |
| Model 11 | $k_D(t,u) = (k_{1,1} \mathbb{1}_{t\leq 24} + k_{2,1} \mathbb{1}_{t>24}) \mathbb{1}_{u<200} + \mathbb{1}_{u\geq 200} (k_{1,2} \mathbb{1}_{t\leq 24} + k_{2,2} \mathbb{1}_{t>24}) e^{\gamma(u-\mu)}$ | $k_{1,1}$ | $2.9 \cdot 10^{-2}$ |
| | | $k_{2,1}$ | $6.4 \cdot 10^{-3}$ |
| | | $k_{1,2}$ | $1.9 \cdot 10^{-2}$ |
| | | $k_{2,2}$ | $9.4 \cdot 10^{3}$ |
| | | $\gamma$ | $1.3 \cdot 10^{3}$ |
| | | $\mu$ | 498 |
| Model 12 | $k_D(t,u) = (k_{1,1} + k_{2,1} e^{-d_1 t}) \mathbb{1}_{u<200} + \mathbb{1}_{u\geq 200} (k_{1,2} + k_{2,2} e^{-d_2 t}) e^{\gamma(u-\mu)}$ | $k_{1,1}$ | $5.2 \cdot 10^{-3}$ |
| | | $k_{2,1}$ | $3.6 \cdot 10^{-2}$ |
| | | $d_1$ | $4.8 \cdot 10^{-1}$ |
| | | $k_{1,2}$ | $5.2 \cdot 10^{-3}$ |
| | | $k_{2,2}$ | $3.6 \cdot 10^{-2}$ |
| | | $d_2$ | $4.7 \cdot 10^{-1}$ |
| | | $\gamma$ | $1.3 \cdot 10^{3}$ |
| | | $\mu$ | 498 |

Table S4: **Parameter values for parametric models of death rate once Smurf.**

| Model | Expression | Parameter value | |
|---|---|---|---|
| Gamma-Gompertz | $k(t) = \frac{\alpha s e^{\alpha t}}{\beta - 1 + e^{\alpha t}}$ | $\alpha$ | $4.0 \cdot 10^{-3}$ |
| | | $s$ | 28.5 |
| | | $\beta$ | 266 |
| Generalized-Gamma | $k(t) = \frac{(p/q)^r t^{r-1} e^{-(t/q)^p}}{\Gamma(r/p) - \gamma(r/p,(t/q)^p}$ | $p$ | 1.5 |
| | | $q$ | 2.2 |
| | | $r$ | 386 |
| Gompertz-Makeham | $k(t) = j + n e^{lt}$ | $j$ | $8.1 \cdot 10^{-4}$ |
| | | $n$ | $9.5 \cdot 10^{-5}$ |
| | | $l$ | $5.8 \cdot 10^{-3}$ |

Table S5: **Parameter values for parametric models of one-phase death hazard rate.**

| Model | Expression | Parameter value | |
|---|---|---|---|
| AKRJ | $k_S(t) = f + g e^{ht}$ | $f$ | $-1.2 \cdot 10^{-3}$ |
| | | $g$ | $1.6 \cdot 10^{-3}$ |
| | | $h$ | $9.8 \cdot 10^{-3}$ |
| | $k_D(t) = k_1 + k_2 e^{-dt}$ | $k_1$ | $6.4 \cdot 10^{-2}$ |
| | | $k_2$ | $-1.4 \cdot 10^{-10}$ |
| | | $d$ | $9.0 \cdot 10^{-3}$ |
| C57 | $k_S(t) = f + g e^{ht}$ | $f$ | $-1.5 \cdot 10^{-3}$ |
| | | $g$ | $1.8 \cdot 10^{-3}$ |
| | | $h$ | $6.7 \cdot 10^{-3}$ |
| | $k_D(t) = k_1 + k_2 e^{-dt}$ | $k_1$ | $3.8 \cdot 10^{-3}$ |
| | | $k_2$ | $8.6 \cdot 10^{-2}$ |
| | | $d$ | $1.9 \cdot 10^{-2}$ |

Table S6: **Parameter values for parametric models of Smurf transition rate and death rate once Smurf for two mice strains (AKRJ and C57).**

| | Model | Parameter value | |
|---|---|---|---|
| DGRP line 377 | Model 6 | $f$ | 0 |
| | | $g$ | $7.5 \cdot 10^{-5}$ |
| | | $h$ | $6.0 \cdot 10^{-3}$ |

Table S7: **Parameter values for parametric two-phase model in DGRP 377 line**. Only $f$, $g$ and $h$ were fit by grid search, the other parameter values used are the same as presented in Table S5.
