## Supplementary Methods for "Acute Smurf mortality and age-dependence in a two-phase ageing model: Statistical inference from drosophila and mice experimental data"

The code, data and Figures can be found at `https://github.com/luce-breuil/smurf_stat`.

#### 1 Data processing

##### 1.1 Drosophila data

The dataset we work with was first published in [13], to which we refer for further details on experimental materials and methods. In brief, mated female flies of synchronised age were maintained individually in vials containing the blue dye, and assessed every 24 hours for life and death as well as the Smurf phenotype. The observations were conducted from day 11 after the birth of the flies until the death of the last fly. For this reason, our study is performed conditionally on the survival of the fly as non-Smurf up to 11 days. In the experimental assay, no fly died before day 11, but some were already Smurf.

The raw dataset is a double entry table with observations made for each individual at different time points. It can be found in the file "smurf.txt". Each entry is an indication on the status of the fly at the considered time (or no indication if the status of the fly has not changed), thus indicating whether the fly is observed Smurf or dead for the first time at that time point. For almost all of the flies, there is only one death indication, but due to experimental uncertainty, a lot of them have several Smurf indications. We chose to keep only the last Smurf observation for each fly, as we consider it to be the most reliable. This is the processing that was done in [13]. The details of the different indications in the raw data are summarized in Table 1 along with the way it is treated.

In general, working with interval censored data adds a layer of complexity and involves iterative methods to compute estimations [18]. For these reasons, we redistribute the jumping times with a uniform distribution within each observation interval. This removes the presence of ties. Note that we also tried a redistribution following a truncated exponential distribution which resulted in more flies transitioning/dying at the beginning of each observation interval, and therefore did not seem likely or adequate. The uniform redistribution we perform is first done on the total lifetimes which are uniformly distributed in the interval in which they were observed. For each fly $i$, we then redistribute the time spent non-Smurf $\tau_i^{NS}$ uniformly in $I \cap \{t \leq \tau_i^D\}$ where $I$ is the 24h interval during which the fly was experimentally observed dying. This ensures that no fly dies before turning Smurf for the flies that were directly observed Smurf and dead. The final distributed times this study was conducted on can be found in jumptimes.txt (for 1159 flies). The values of the parameters fitted for the Smurf transition rate are in Table S4, for the death rate in Table S5 and for one-phase death hazard rates in Table S6.

In this study, we work only with the uniformly distributed data but some of the nonparametric estimations were also conducted on the exponentially distributed data and the censored data as well and the conclusions were unchanged.

| symbol | meaning | treatment |
|---|---|---|
| s | smurf | kept |
| sl | smurf light | replaced by s |
| ns | non smurf after being marked s? or s | removed (228 indications) |
| sd | smurf dead | kept |
| sld | smurf light dead | replaced by sd |
| lost/perdue | lost data or fly | line removed (6 flies) |
| dns | dead non smurf: fly found dead attached to the side of the tube, could not access the blue coloring | line removed (1 fly) |
| s? | not sure if smurf or not | removed (359) |
| sd from beginning | already smurf and dead when experiment begins | removed (30 flies) |
| sl after sd | experimental error | removed (1 fly ) |
| fly not dead during trial | lost data | line removed (3 flies) |
| fly turned Smurf within first 11 days | fly turned smurf in first 11 days | line removed (41 flies) |

Table 1: Symbols for the experimental observations, their meaning and treatment

Another Drosophila dataset was used for Figure S9. These flies are from the Drosophila Genetic Resource Panel (DGRP) line 377 and their Smurfness and survival was assessed while being kept in vials of $\sim 30$ individuals as described in [13]. For this reason, the data only consists in population survival data and a few time points for which the Smurf proportion is observed. On this data, only parameters $g$ and $h$ were fitted by maximizing the likelihood of the law of total lifetimes on a grid search. The parameter values can be found in Table S8.

#### 1.2 Mice data

The dataset of mice survival is taken from [3]. It consists in the observation of time spent non-Smurf and Smurf of mice from two different strains (AKRJ female and C57 male and female). The mice were tested for several indicators such as inflammation, metabolic parameters, microbiota and intestinal permeability at each observation time. In [3], they determine the point at which each mouse turns Smurf by finding the moment when the health indicators concurrently fail. We refer the reader to [3] for more details on the data. Due to the large observation interval of 3 weeks, several mice were directly observed dead without turning Smurf beforehand. As our model assumes a 2-phase aging, we distribute the time spent Smurf of these mice within the observation interval prior to their observed death. Unlike what was done for the Drosophila dataset, we do not uniformly distribute the observations. The treatment of the data can be found in the files Mice_treatment.R and Mice.ipynb. After treatment, the dataset for the AKRJ mice strain is of size 45 and for the C57 mice of size 92. The values of the fitted parameters for both mice strains can be found in Table S7.

#### 2 Statistical estimation

Let $\tau$ be an absolutely continuous positive random variable (i.e. a random variable which admits a probability density function). The law of $\tau$ can alternatively be described by its distribution function $F(t) = \mathbb{P}(\tau \leq t)$, its density $f(t) = \frac{\mathrm{d}F}{\mathrm{d}t}$ or its hazard rate $k$ defined by

$$k(t) = \frac{f(t)}{1 - F(t)}, \tag{1}$$

where the quantity $1 - F$ is called the survival function associated to the random variable. The hazard rate $k(t)$ can be interpreted as the probability that an individual experiences an event during an infinitesimal time interval $(t, t + \mathrm{d}t)$, conditionally on having survived until time $t$. In survival analysis and demography, the hazard rate can be referred to as force of mortality or mortality rate. In our case, we are interested in estimating the hazard rate $k$ of a random variable $\tau$ with an observed sample of size $m$, $(\tau_i)_{1 \leq i \leq m}$.

##### 2.1 Nonparametric estimation

When the shape of the hazard rate function is completely unknown, parametric approaches are not well suited and can result in biased estimations. In such cases, nonparametric approaches can be particularly useful for estimating the hazard rate without prior knowledge. Kernel estimators are among the most widely used nonparametric estimators. They were first introduced and used for density estimation [10] and for hazard rate estimation in [15, 16]. In order to estimate the hazard rate $k$ associated to an observed series of times $(\tau_i)_{1 \leq i \leq m}$, we use a nonparametric estimation with the Gamma kernel $\kappa_{t,b}$ as described in [2]. For $t \geq 0$ and $b > 0$, the Gamma kernel at point $t$ of bandwidth $b$ is defined by

$$\kappa_{t,b}(y) = \frac{y^{\rho(t)_b - 1} e^{-y/b}}{b^{\rho(t)_b} \Gamma(\rho(t)_b)} \mathbb{1}_{\{y \geq 0\}} \tag{2}$$

where

$$\rho(t)_b = \begin{cases} t/b \text{ if } t \geq 2b \\ \frac{1}{4}(t/b)^2 + 1 \text{ if } 0 \leq t < 2b. \end{cases} \tag{3}$$

The hazard rate kernel estimator at point $t$, $\hat{k}(t)$ is defined as follows

$$\hat{k}(t) = \sum_{i \geq 1} \frac{1}{m - N_{\tau_i^-}} \kappa_{t,b}(\tau_i) \text{ where } N_t = \sum_{i=1}^{m} \mathbb{1}_{\{\tau_i \leq t\}}, \quad \forall t \geq 0. \tag{4}$$

The bandwidth $b$ is chosen with the local minimax bandwidth choice presented in Section 4.2 of [2].

To estimate the cumulated base hazard in a Cox model of dependence, we use the Breslow estimator, which is described in [7]. It consists in estimating the base hazard with a piecewise constant function, taking constant values between observed jumping times. The values of the constants are determined by maximum likelihood. The implementation of the estimator we use is in the function `coxph` of the package survival in R.

##### 2.2 Maximum likelihood parametric estimation

In the case where the shape of a function is known, parametric estimation allows for an interpretable and comparable way of estimating the function. It also results in an explicit expression of the function. In our case, this approach makes it possible to compare models quantitatively among each other, and with other classical models found in the literature. In order to determine the parameters in our parametric estimations, we use the maximum likelihood estimator for the rate of jump processes as described in Chapter VI in [1].

We set $\tau_0 = 0$ and for $t \geq 0$, $N_t = \sum_{i=1}^{m} \mathbb{1}_{\{t \geq \tau_i\}}$, the counting process associated to the times $(\tau_i)_{1 \leq i \leq m}$. $N_t$ represents the number of events (in our case, transitions or deaths) having occurred before time $t$. Let $\bar{k}(t,\theta)$ be a parametric function of $t$ and $\theta = (\theta_1, ..., \theta_l)$ the parameters we wish to determine. $\bar{k}(t,\theta)$ is a parametric estimation of $k$.

For a given parametrisation $\bar{k}$, the log-likelihood is

$$L(\bar{k},\theta) = \int_0^{\tau_m} \log(\bar{k}(t,\theta)N_s)dN_s - \int_0^{\tau_m} \bar{k}(t,\theta)N_s\,\mathrm{d}s$$
$$= \sum_{k=1}^{m} \log(\bar{k}(\tau_k,\theta)) + \sum_{k=1}^{m} \log(N_{\tau_k}) - \int_0^{\tau_m} \bar{k}(t,\theta)N_s\,\mathrm{d}s. \tag{5}$$

Thus by differentiating (5) with respect to each parameter $\theta_j$, the maximum likelihood parameters solve the following system

$$\sum_{k=1}^{m} \frac{\partial \bar{k}(\tau_k,\theta)}{\partial \theta_j} \frac{1}{\bar{k}(\tau_k,\theta)} - \int_0^{\tau_m} \frac{\partial \bar{k}(t,\theta)}{\partial \theta_j} N_s\,\mathrm{d}s = 0 \text{ for } j = 1, ...l. \tag{6}$$

Since $N_s$ is piecewise constant with $N_s = N_{\tau_k}$ on $[\tau_k, \tau_{k+1}]$, (6) simplifies to

$$\sum_{k=1}^{m} \frac{\partial \bar{k}(\tau_k,\theta)}{\partial \theta_j} \frac{1}{\bar{k}(\tau_k,\theta)} - \sum_{k=1}^{m} N_{\tau_{k-1}} \int_{\tau_{k-1}}^{\tau_k} \frac{\partial \bar{k}(t,\theta)}{\partial \theta_j}\,\mathrm{d}s = 0 \text{ for } j = 1, ...l. \tag{7}$$

Solving (7), and checking that the parameter values obtained are indeed maximizers of the log-likelihood, allows to find the maximum likelihood estimator $\hat{\theta}$ of $\theta$.

95% confidence intervals for the maximum likelihood estimators of the parameters can be obtained. We first compute the Fischer Information, which is an $l \times l$ matrix defined by (see e.g. [17, 8])

$$I_m(\hat{\theta}) = \left( -\mathbb{E}\left[ \frac{\partial^2}{\partial \theta_i \partial \theta_j} \log(\bar{k}(t,\hat{\theta}) e^{-\int_0^{\tau_m} \bar{k}(s,\hat{\theta})\,\mathrm{d}u}) \right] \right)_{i,j \in [1,l]}. \tag{8}$$

Let $A_m(\hat{\theta}) = \sqrt{I_m(\hat{\theta})^{-1}}$ be the matrix such that $A_m(\hat{\theta})^2 = I_m(\hat{\theta})^{-1}$. The 95% confidence interval for each component $\hat{\theta}_i$ of $\hat{\theta}$ is defined by

$$CI(\hat{\theta}_i) = \left[ \hat{\theta}_i \pm \frac{q_\alpha}{\sqrt{m}} A_{m,i,i}(\hat{\theta}) \right]. \tag{9}$$

#### 2.3 Cox Model of dependence

We model the dependence between $\tau^S$ and $\tau^{NS}$ with Cox's proportional hazard model [6] which is common in survival analysis to model the impact of a covariable on the hazard rate of an event. This model hypothesises an exponential dependence between the hazard rate and the covariable, in our case, between the death hazard rate of an individual and the time it spent non-Smurf $\tau^{NS}$. The most general form for the death hazard rate $k_D$ that we consider is

$$k_D(a,v) = \sum_{j=1}^{m_0} \mathbb{1}_{\{v \in I_j\}} k_{D,j}(a) e^{\gamma g(v)} \tag{10}$$

where $a$ is the Smurf age and $v$ the covariable, $g$ is a function and $\{I_j,\ j \in [1, m_0]\}$ is a set of intervals. $\gamma$ is called the Cox coefficient and if $\gamma = 0$, there is no dependence on the covariable $v$ ($\tau^{NS}$ in our case). We refer to the functions $k_{D,j}(a)$ as the base hazards, which can be different on different intervals of $v$ (this corresponds to the stratified Cox model [5]), or equal, which is the case in the classical Cox model.

To estimate the parameter $\gamma$, we reorder the $(\tau^{NS})_{1\leq i\leq m}$ such that $\tau_{i-1}^{NS} \leq \tau_i^{NS} \leq \tau_{i+1}^{NS}$. The parameter $\gamma$ is estimated by maximizing the following log partial likelihood function defined in [12]

$$l(\gamma) = \sum_{i=1}^{m} \left( \gamma g(\tau_i^{NS}) - \log \left( \sum_{\substack{r\in[\![1,m]\!] \\ \tau_r^S \geq \tau_i^S}} e^{\gamma g(\tau_r^{NS})} \right) \right). \tag{11}$$

Notice that there is no dependence on the base hazard in (11), meaning that $\gamma$ and the base hazard functions are fitted and studied independently. Once the Cox coefficient is determined by maximizing (11), the parameters of the base hazard can be fit by maximum likelihood by solving (7). The Wald test of independence quantifies how significantly $\gamma$ is different from 0 in a Cox model of dependence.

### 3 Models comparison

The indicators we use to compare the models are as follows

- **The Bayesian Information criterion (BIC)** [9], which can be interpreted as an approximation of the Bayesian posterior probability of a model. For a parametrisation $\bar{k}$ with parameters $\hat{\theta}$, the BIC is defined by

$$\text{BIC}(\bar{k}, \hat{\theta}) = -2 \log L(\bar{k}, \hat{\theta}) + l(\bar{k}) \log(m) \tag{12}$$

  with $l(\bar{k})$ the number of fitted parameters, which depends on the chosen parametrisation, and $m$ the sample size. $L(\bar{k}, \hat{\theta})$ is the log-likelihood of the model with hazard rate $\bar{k}$ and parameters $\hat{\theta}$. This criterion comes down to penalizing models that have a lot of parameters to prevent overfitting. Furthermore, the BIC is asymptotically consistent as the sample size grows, meaning that the penalty divided by the sample size converges to zero [14]. This means that if the true model is part of the tested models, the $BIC$ will select it as the best one when the sample size grows to infinity. Better models are thus assumed to minimise the BIC. For a fixed sample size however, it is only a heuristics and does not constitute a proof that a parametrisation is better than another one [9].

- **The empirical Wasserstein distance** between the distribution of the data and the model distribution [4]. Let $(X_i)_{(1\leq i\leq m)}$ be experimental data of empirical law $\hat{\mu}$ and $(Y_i)_{(1\leq i\leq m)}$ be the corresponding simulated data of empirical law $\hat{\nu}$. The empirical Wasserstein distance is defined as follows

$$W_m(\hat{\mu}, \hat{\nu}) = \inf_{\sigma\in S_m} \frac{1}{m} \sum_{i=1}^{m} ||X_i - Y_{\sigma(i)}||. \tag{13}$$

  where $S_m$ denotes the set of permutations of $[\![1, m]\!]$. Depending on whether we are looking at the law of one transition time only or of the joint law of both transition times, the norm $||.||$ considered is the absolute value (1D) or the euclidian norm in 2-dimensions $d((x_1, x_2), (y_1, y_2)) = \sqrt{(x_1 - y_1)^2 + (x_1 - y_2)^2}$ (2D). To compute the Wasserstein distances, we use the functions `wasserstein` and `wasserstein1D` of the package transport in R [11]. As the empirical Wasserstein distance between our data and one simulation of a model is highly variable, we consider the mean empirical distance between the data and 100 simulations of a given model. It is worth mentioning that the Wasserstein distance does not penalise models with a lot of parameters thus not accounting for overfitting (unlike the BIC).

- **QQ-plots**, or quantile-quantile plots, are figures where the quantiles of a theoretical simulated distribution are plotted against the quantiles of the empirical distribution of data. In our case, the quantiles are computed on a simulated population of 12000 individuals. If both distributions are equal, the quantiles will be equal and the plotted points will align on the $x = y$ line. The visual distance from the $x = y$ line is therefore an indication of how close the distributions are to each other.

- **95% error bands/Confidence intervals**. The error bands shown in several of our figures are done by simulating 500 populations and plotting pointwise the 2.5% and 97.5% quantiles of the quantity of interest over the 500 simulations. The confidence intervals shown (for parametric and nonparametric estimations), are computed using either equation (9) for maximum likelihood parametric estimations or Theorem 3.2 in [2] with the variance and expectation of the estimator estimated by Monte-Carlo for the nonparametric estimation. As the number of individuals decays with time, the confidence intervals widen, showing the lack of confidence we expect when only few individuals remain alive at the time of estimation.
- **Visual comparison of survival function** The use of the non-Smurf survival function as an indicator of goodness-of-fit is done since it is a quantity that can be represented empirically from the data without preprocessing, contrarily to what would be necessary for the density and the transition rate.
